# Deep learning for histopathological subtyping and grading of lung adenocarcinoma

**DOI:** 10.1101/2022.11.02.514811

**Authors:** Kris Lami, Noriaki Ota, Shinsuke Yamaoka, Andrey Bychkov, Keitaro Matsumoto, Wataru Uegami, Richard Attanoos, Sabina Berezowska, Luka Brcic, Alberto Cavazza, John C. English, Alexandre Todorovic Fabro, Kaori Ishida, Yukio Kashima, Yuka Kitamura, Brandon T. Larsen, Alberto M. Marchevsky, Takuro Miyazaki, Shimpei Morimoto, Mutsumi Ozasa, Anja C. Roden, Frank Schneider, Maxwell L. Smith, Kazuhiro Tabata, Angela M. Takano, Tomonori Tanaka, Tomoshi Tsuchiya, Takeshi Nagayasu, Hidenori Sakanashi, Junya Fukuoka

## Abstract

The histopathological distinction of lung adenocarcinoma (LADC) subtypes is subject to high inter-observer variability, which can compromise the optimal assessment of the patient prognosis. Therefore, this study developed convolutional neural networks (CNNs) capable of distinguishing LADC subtypes and predicting disease-specific survival, according to the LADC tumour grades established recently by the International Association for the Study of Lung Cancer pathology committee. Consensus LADC ground truth histopathological images were obtained from seventeen expert pulmonary pathologists and one pathologist in training. Two deep learning models (AI-1 and AI-2) were trained with EfficientNet b3 architecture to predict eight different LADC classes (lepidic, acinar, papillary, micropapillary, solid, invasive mucinous adenocarcinoma, other carcinoma types, and no carcinoma cells). Furthermore, the trained models were tested on an independent cohort of 133 patients. The models achieved high precision, recall, and F1-scores exceeding 0.90 for most of the LADC classes. Clear stratification of the three LADC grades was reached in predicting the disease-specific survival by the two models. Moreover, the grading prediction of one of the trained models was more accurate than those of 14 out of 15 pulmonary pathologists involved in the study (*p*=0.0003). Both trained models showed high stability in the segmentation of each pair of predicted grades with low variation in the hazard ratio across 200 bootstrapped samples. These findings indicate that the trained CNNs improve the diagnostic accuracy of the pathologist, standardise LADC subtype recognition, and refine LADC grade assessment. Thus, the trained models are promising tools that may assist in the routine evaluation of LADC subtypes and grades in clinical practice.

## INTRODUCTION

In recent years, advances in digital pathology with the use of whole slide images (WSIs) have enabled the introduction of machine learning in the pathology domain, and deep learning algorithms as well as computer-aided diagnostic tools have been established [1–3]. Numerous deep learning models have been developed in the area of diagnostic pathology for breast cancer [4–6], prostate cancer [7–9], soft tissue sarcomas [10], hematologic malignancies [11], etc. Moreover, in lung pathology, several convolutional neural network (CNN) models have been built for various tasks [12–15], including the distinction of the different histologic types of lung cancer [16–18].

Lung adenocarcinoma (LADC), which is a subtype of non-small cell lung cancer, is the most common histologic type of lung cancer across the globe [19]. It is a heterogeneous entity subdivided into adenocarcinoma in situ, minimally invasive adenocarcinoma, invasive non-mucinous adenocarcinoma, invasive mucinous adenocarcinoma (IMA), colloid adenocarcinoma, foetal adenocarcinoma, and enteric-type adenocarcinoma. Invasive non-mucinous adenocarcinoma, in turn, comprises several subtypes according to architectural patterns: lepidic (a non-invasive pattern), acinar, papillary, micropapillary, and solid [20]. The distinction of these histologic subtypes is critical because some of them are independent prognostic factors. For instance, the poor prognosis when micropapillary and solid subtypes are present in a tumour is well-documented, even as non-predominant subtypes [21–26]. It has been demonstrated that cribriform arrangements of tumour glands or complex glandular patterns, described within the acinar subtype of LADC, exhibit aggressive behaviour with poor prognosis [27,28]. Semiquantitative reporting of these subtypes is recommended in 5% increments to include even a small amount of prognostically significant subtypes [29]. However, several studies have indicated that the distinction of the different LADC subtypes is subjective and prone to high inter-observer variability [30–32]. Our previous research showed an overall fair agreement (κ = 0.338) between pulmonary pathologists when determining the predominant LADC subtype in a series of cases, with an agreement for the distinction of specific predominant subtypes varying from slight to substantial [33]. Several CNNs have been developed for assisting pathologists in recognising and quantifying the LADC subtypes [34,35]. However, not all LADC subtypes have been included in these studies (IMA for example, which accounts for 3–10% of LADC [20], was not included in these models). Furthermore, with known interobserver variability, the ground truth of the LADC growth patterns was not always guaranteed as it was obtained by not more than 3 pathologists.

In this study, we attempted to train and develop CNN models capable of distinguishing the LADC subtypes, based on a set of consensus LADC images provided by 17 international expert pulmonary pathologists and one pathologist in training obtained through a clustering approach in our previous study [33]. Furthermore, the trained algorithms predicted survival based on the novel LADC grading system proposed by the International Association for the Study of Lung Cancer (IASLC) pathology committee [36].

## MATERIALS AND METHODS

### Study design and data collection

A retrospective study, approved by the Clinical Research Review Committee of the Nagasaki University Hospital (no. 20042008-2), was carried out. A cohort of 191 surgically resected primary LADC cases from the Nagasaki University Hospital (Nagasaki, Japan) was selected. After retrieving and scanning the glass slides at 20x magnification (Aperio Scanscope CS2 digital slide scanner, Leica Biosystems, Buffalo Grove, USA), 330 WSIs were obtained. They were further randomly divided into a training set and a test set. In total, 91 and 239 WSIs were used for model training and testing, respectively.

### Labelling (annotation) of the training set

From the 91 WSIs of the training set, 12 WSIs comprising each subtype of LADC were selected and segmented into 1 mm^2^ tile with a resolution of 2 μm per pixel. After excluding tiles with more than 80% blank background, 4,702 tiles were obtained. As described in our previous study [33], 17 expert pulmonary pathologists and one pathologist in training from 16 institutions representing 9 countries labelled each tile according to the predominant subtype present. These labels included lepidic, acinar, papillary, micropapillary, solid, and IMA. “Other carcinoma types” was included in the labels to represent cribriform and complex glandular architecture, as well as other rare LADC subtypes. Tiles devoid of tumour cells, containing benign lung parenchyma or other non-neoplastic features were labelled as “No carcinoma cells”.

The relatively uniform tumour architectural areas of the remaining 79 WSIs were annotated (ASAP, Computation Pathology Group, Nijmegen, The Netherlands), and the areas were labelled as a LADC subtype by each pathologist. After labelling, each annotated area was segmented in a similar manner as the previous 12 WSIs. Including tiles containing more than 25% of tissue only, 8,554 tumour tiles were obtained. In addition to the uniform cancerous areas, the benign areas of the 79 WSIs were annotated and segmented. After excluding tiles containing less than 50% of tissue, segmentation resulted in 19,204 non-cancerous tiles.

### Ground truth for the training set

The clustering approach used for obtaining a reliable set of ground truth images for the LADC subtypes is described in our previous study [33]. The cluster analysis was used to highlight pathologists with similar morphological pattern recognition. Two clusters of pathologists were formed, based on the labels of the 4,702 tiles. The main difference between the clusters was the assessment of certain patterns such as IMA by Cluster-1, and micropapillary or other cancer types by Cluster-2. In Cluster-1 containing 10 pathologists, consensus for a tile was set as 6-out-of-10 identical LADC subtype labels. In Cluster-2 containing 5 pathologists, the consensus was set as 3-out-of-5 identical labels. Three pathologists were outliers. Thereby, among the 4,702 tiles from the 12 WSIs, 3,733 tiles had consensus in Cluster-1 and 4,214 tiles in Cluster-2. Of the 8,554 tumour tiles originating from the 79 WSIs, 6,409 tiles had consensus in Cluster-1 and 7,188 in Cluster-2. As a result, a total of 10,142 and 11,402 consensus tiles of different LADC subtypes were obtained for Cluster-1 and Cluster-2, respectively (Figure 1).

**Figure 1.**
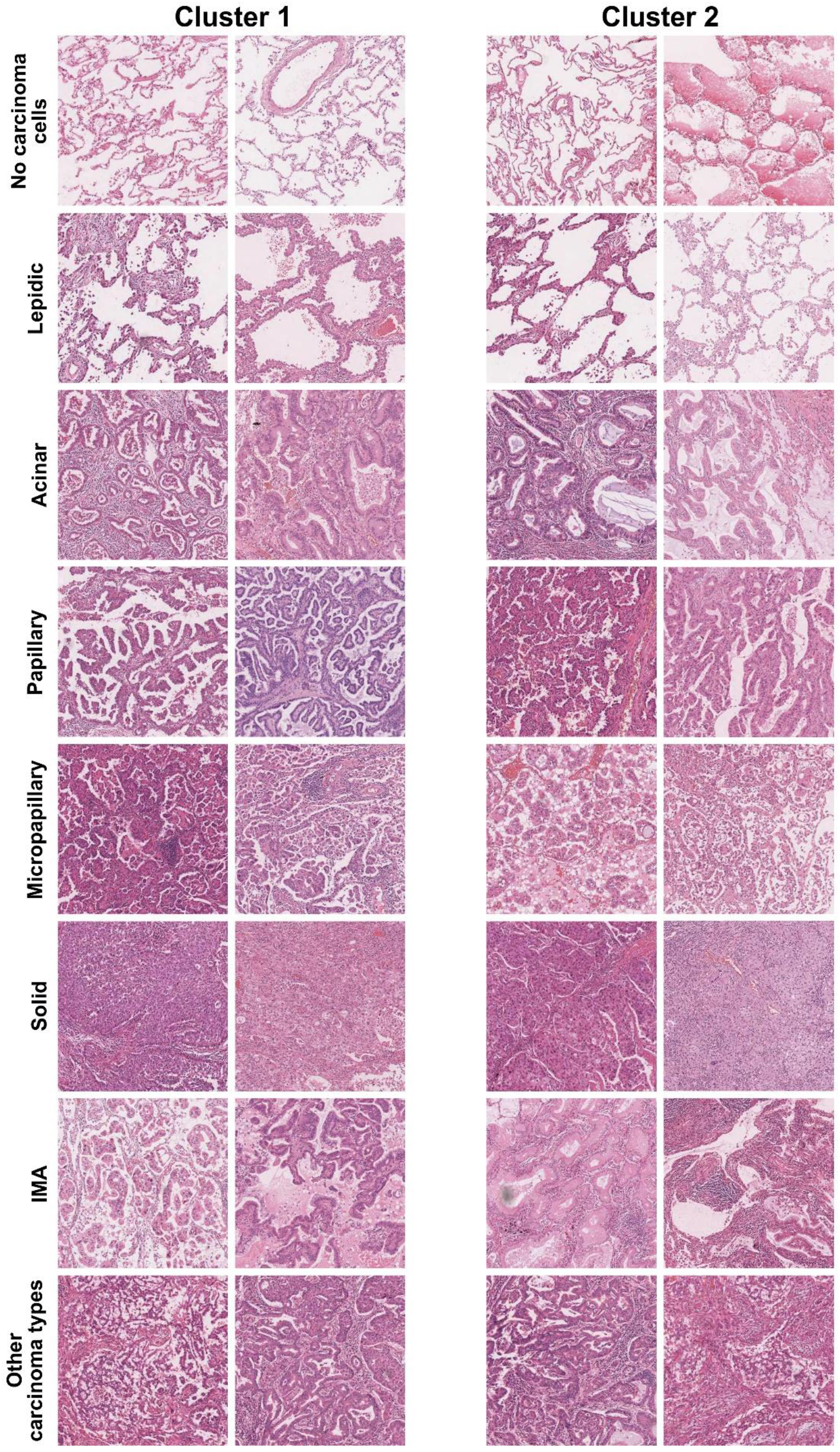
Representative tiles of the lung adenocarcinoma histopathological classes used for models training IMA, invasive mucinous adenocarcinoma

### Training of the deep learning models

Two models, AI-1 and AI-2, were trained in this study. The training set of AI-1 comprised the 10,142 consensus tiles of Cluster-1 and 19,204 non-cancerous tiles, resulting in a total of 29,346 tiles. The training set of AI-2 comprised the 11,402 consensus tiles from Cluster-2 and 19,204 non-cancerous tiles, resulting in 30,606 tiles (Figure 2A).

**Figure 2.**
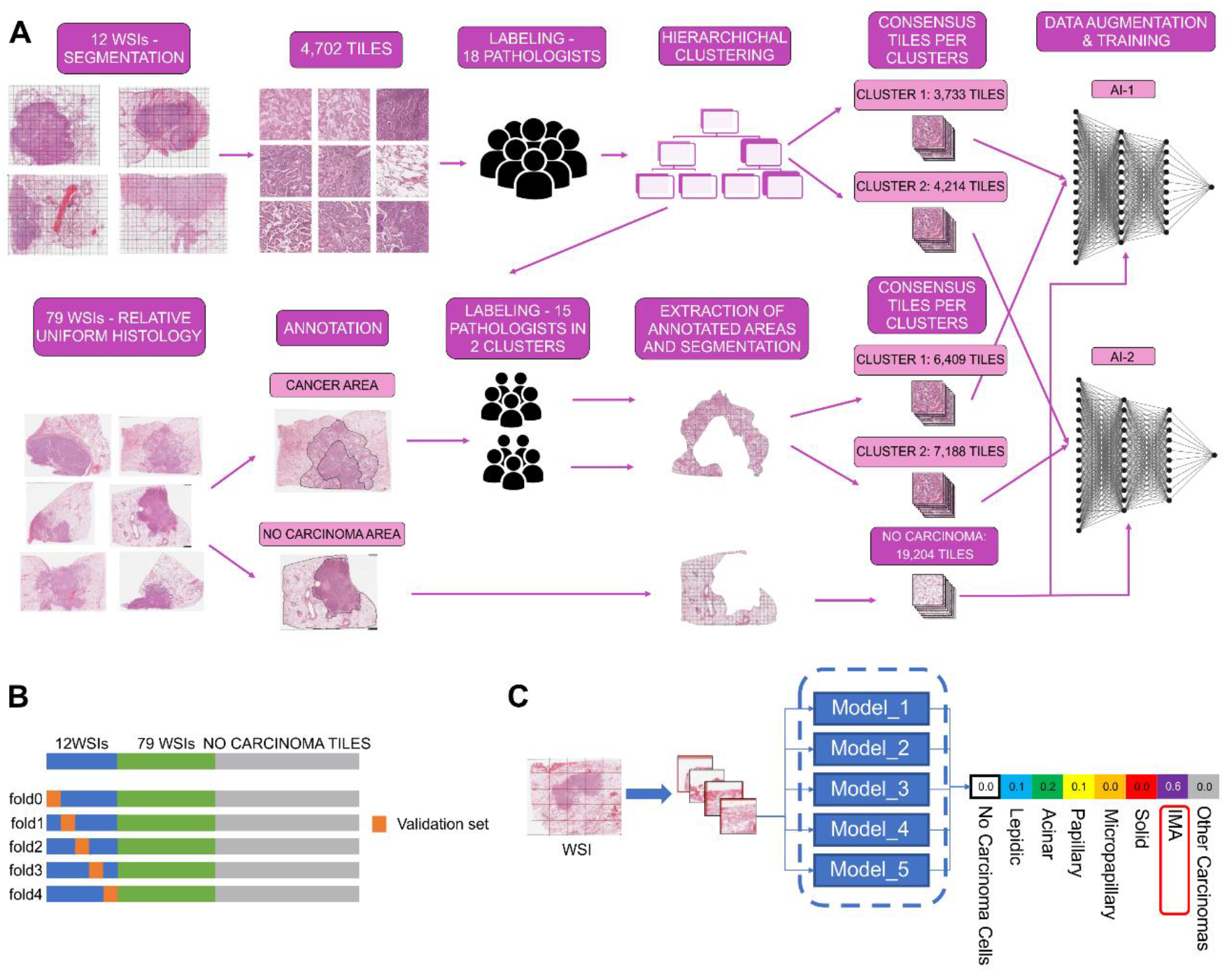
A pipeline of deep learning model development. (A) Model training strategies. Two models (AI-1 and AI-2) are trained based on the consensus labels obtained from the initial 18 pathologists through a hierarchical clustering approach. (B) Model validation procedure. A five-fold cross-validation method is applied to validate the models, with a set of 91 WSIs. (C) Model testing. Based on the averaged results of the five-fold cross-validation, the final inference result with the highest prediction score is selected from the class.

The two models were trained with the EfficientNet-B3 architecture, using the timm library [37]; tf_efficientnet_b3_ns was used as a pre-trained parameter. The input image size was 512 x 512 with a batch size of 64. The two models learned a total of 100 Epochs. Adam was employed as the optimiser. The initial learning rate was 0.001, which was attenuated for each epoch based on cosine annealing. Binary cross entropy was applied as the loss function. The training was performed using single hardware (NVIDIA V100 32 GB GPU, Nvidia Corporation, Santa Clara, USA), with mixed-precision training to speed up the training process.

### Data augmentation

Each tile was loaded and augmented immediately before being put into the models. Pixel-level and spatial-level transforms from the Albumentations library [38] and the mixup principle [39] were applied to each tile (supplementary material, Figure S1). Details on how the transforms were applied are shown in Figure 3.

**Figure 3.**
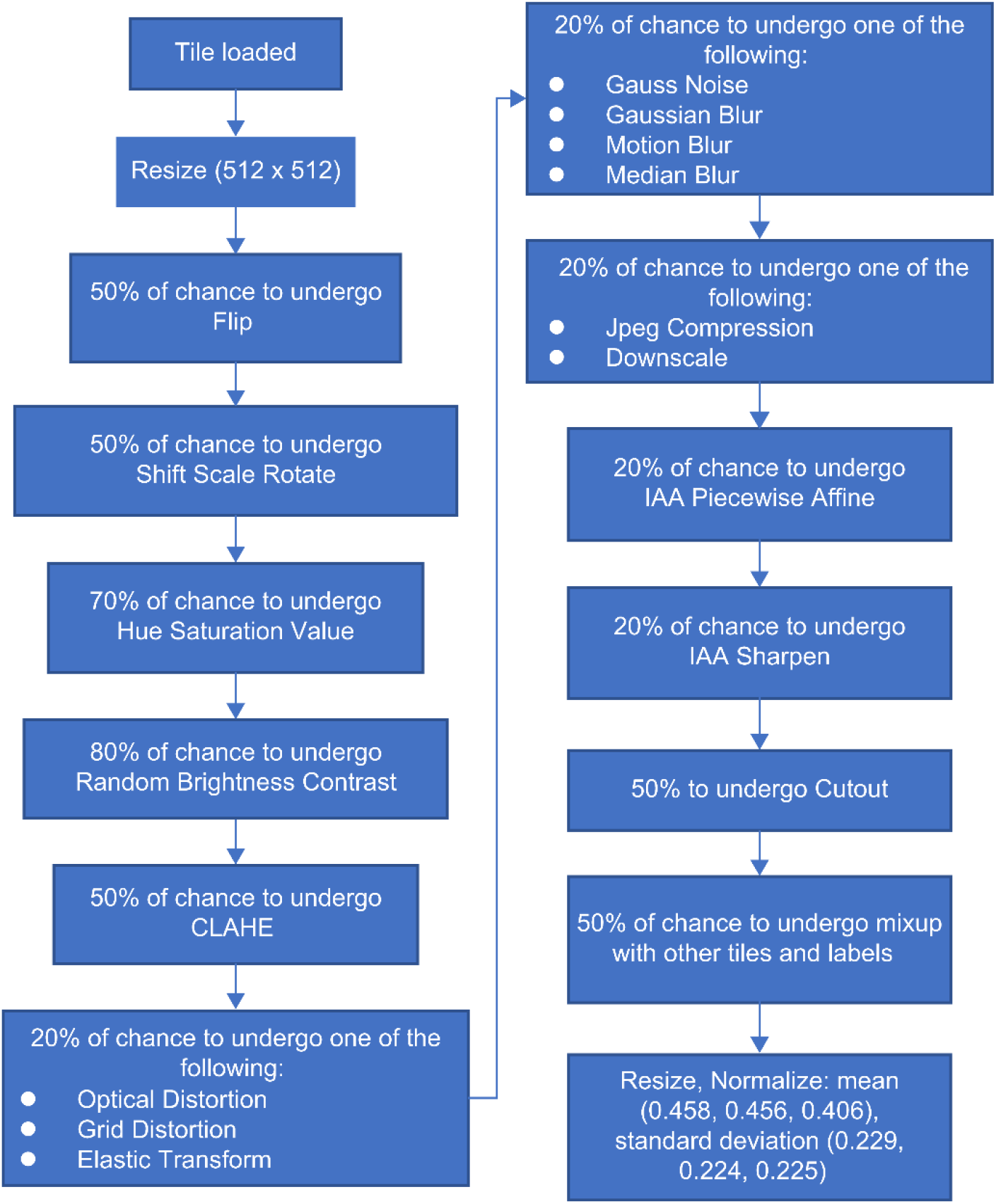
Application of Albumentation transforms and mixup principle on a tile

### Validation of models

The k-fold cross-validation process (k = 5) was applied for validating the trained models. The tiles obtained from the 12 WSIs were divided into training (80%) and validation (20%) sets. The distribution was performed in a way that a tile obtained from a WSI was not included in the training and validation sets at the same time, and homogeneity was ensured between the training and validation sets in terms of the LADC subtypes. All the tiles from the 79 WSIs were included in the training set (tumour and non-tumour tiles) (Figure 2B). The models were validated against the consensus diagnoses of each tile of Cluster-1 for AI-1 and Cluster-2 for AI-2. The validation score was checked for each epoch, and the checkpoint with the highest validation accuracy was used as the folding model for AI-1 and AI-2.

### Labelling of the test set

The remaining fifteen pathologists provided case-level diagnoses of the 142 cases from the 239 WSIs. Each LADC subtype was recorded with an estimation of the percentage in 5% increments. Further, the tumour grades were evaluated for each case considering the predominant patterns and high-grade patterns [36]. Since the grading system was not validated in variants of LADC, cases labelled as IMA-predominant tumour by at least one of the 15 pathologists were excluded, resulting in 133 cases with LADC grades.

### Models testing

Each WSI from the test set was segmented into tiles and pre-processed with the same parameters as the tiles originating from the WSIs of the training set (1 mm^2^ tile with a per-pixel resolution of 2 μm; resizing and normalisation). LADC class inferences of the trained models were obtained using the five models trained through cross-validation on the pre-processed patch image. The inferred results were averaged, and the class (lepidic, acinar, papillary, micropapillary, solid, IMA, other carcinoma types, or no carcinoma cells) with the highest score was adopted as the inference result (Figure 2C). An inference map was drawn using the tiles with their respective inference results highlighted by colour codes and superimposed on the corresponding original WSI. The percentages of all the LADC subtypes were then calculated and the LADC grades were evaluated for each case.

### Statistical analysis

The precision, recall, and F1 score of each class by the trained models were calculated using the consensus labels of the respective clusters as the ground truth. Confidence intervals of 95% were obtained based on the bootstrap method by iterating the results of cross-validation 1000 times. The receiver operating characteristic (ROC) curves and the resulting area under the curve (AUC) values were calculated and plotted using Python (version 3.7.7), with the scikit-learn (version 1.0.1) and Matplotlib (version 3.3.4) packages. The pairwise agreements between the pathologists and trained models were evaluated based on Cohen’s kappa coefficient. The disease-specific survival (DSS) of the test set based on the LADC grades estimated by the pathologists and the trained models were calculated using the Kaplan-Meier estimator in JMP Pro, version 15.0.0 (SAS Institute, Cary, North Carolina, USA). The survival curves were judged statistically significant if the *p* values were less than 0.05.

For each of the pathologists and trained models, the grading performance as the sensitivity in the segmentation of the cases of the test set, from the point of view of the survival rate, was evaluated based on the hazard ratio between the LADC grades diagnosed by the pathologist or predicted by the trained models (grade 1 versus grade 2; grade 1 versus grade 3; grade 2 versus grade 3). The rank and stability of the performance were evaluated based on the median and range (25^th^ – 75^th^ percentile) of the estimated hazard ratio in 200 trials of bootstrap sampling. The hazard ratio was estimated by the Cox proportional hazard model.

## RESULTS

### Cross-Validation

Consensus tiles originating from the whole slide segmentation of 12 WSIs and the uniform area segmentation of 79 WSIs were used for training the models. Additionally, the non-cancerous tiles from the same 79 WSIs were added to the training set. The number of training tiles per LADC subtype per model is shown in Table 1.

**Table 1:** Number of training tiles per lung adenocarcinoma subtype per model.

|  | **AI-1** | **AI-2** |
| --- | --- | --- |
| **Subtypes** |  |  |
| No carcinoma cells | 21,426 | 21,380 |
| Lepidic | 860 | 1,170 |
| Acinar | 1,861 | 2,399 |
| Papillary | 1,591 | 2,220 |
| Micropapillary | 608 | 741 |
| Solid | 1,185 | 1,405 |
| IMA | 1,781 | 1,207 |
| Other carcinoma types | 34 | 84 |

The two trained models (AI-1 and AI-2) were validated through five-fold cross-validation. The number of tiles used for validating each fold is listed in Table 2.

**Table 2:** Cross-validation characteristics.

|  | Number of training tiles | Fold | Number of validation tiles | Batch size | Epoch | Number of iterations |
| --- | --- | --- | --- | --- | --- | --- |
| **Models** |  |  |  |  |  |  |
| **AI-1** | 29,346 | 0 | 977 | 64 | 100 | 44,300 |
|  |  | 1 | 213 |  |  | 45,500 |
|  |  | 2 | 890 |  |  | 44,400 |
|  |  | 3 | 881 |  |  | 44,400 |
|  |  | 4 | 772 |  |  | 44,600 |
| **AI-2** | 30,606 | 0 | 671 | 64 | 100 | 46,700 |
|  |  | 1 | 1070 |  |  | 46,100 |
|  |  | 2 | 908 |  |  | 46,400 |
|  |  | 3 | 616 |  |  | 46,800 |
|  |  | 4 | 949 |  |  | 46,300 |

The confusion matrices showing the performances of the trained models in predicting the LADC subtypes against the consensus obtained from the respective clusters are displayed in Figures 4A and 4B, respectively.

**Figure 4.**
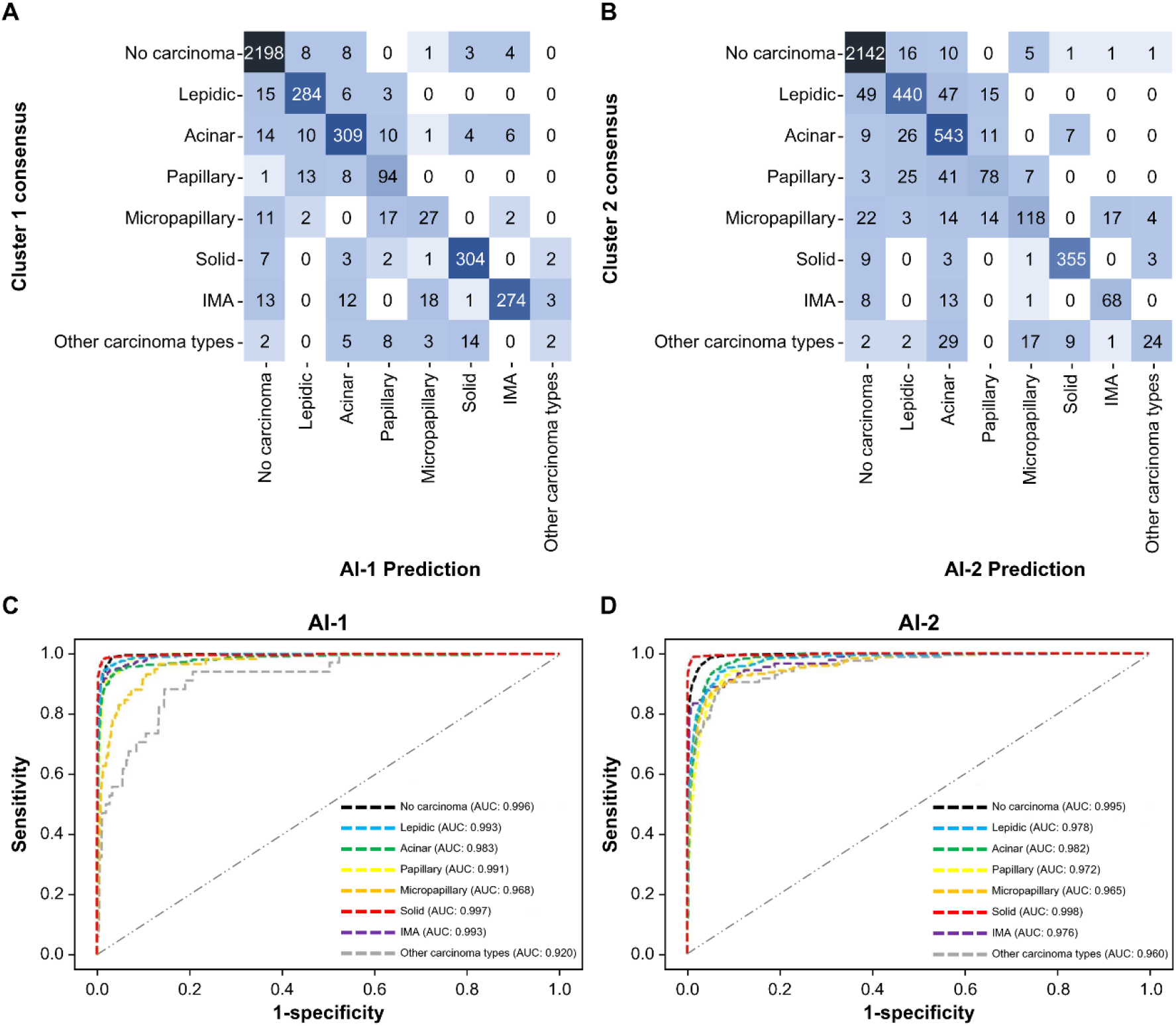
Cross-validation results. (A, B) Confusion matrices of the trained-model predictions against the corresponding cluster consensus. (C and 3D) Receiver operating characteristic (ROC) curves for each predicted class with the corresponding AUCs. Examples of tiles with various degrees of agreement are shown in supplementary material, Figure S2 IMA, invasive mucinous adenocarcinoma; AUC, area under the curve

Both AI-1 and AI-2 showed high performance in predicting the LADC classes with accuracies above 97% and 95%, respectively. AI-1 achieved a precision of more than 0.90 for the “no carcinoma”, “IMA”, and “solid” classes (0.972, 0.958, and 0.933, respectively), whereas AI-2 showed similar results for the “no carcinoma” and “solid” classes (0.954 and 0.955, respectively) (Table 3). The recall score exceeded 0.90 for the “no carcinoma”, “lepidic”, and “solid” classes (0.989, 0.921, and 0.953, respectively) with AI-1, whereas AI-2 achieved a comparable score for the “no carcinoma”, “acinar”, and “solid” classes (0.984, 0.911, and 0.956, respectively). The “no carcinoma” and “solid” classes obtained the highest F1 score among all the classes for both AI-1 and AI-2 (0.980 and 0.943 for AI1, 0.970 and 0.956 for AI-2).

**Table 3:**
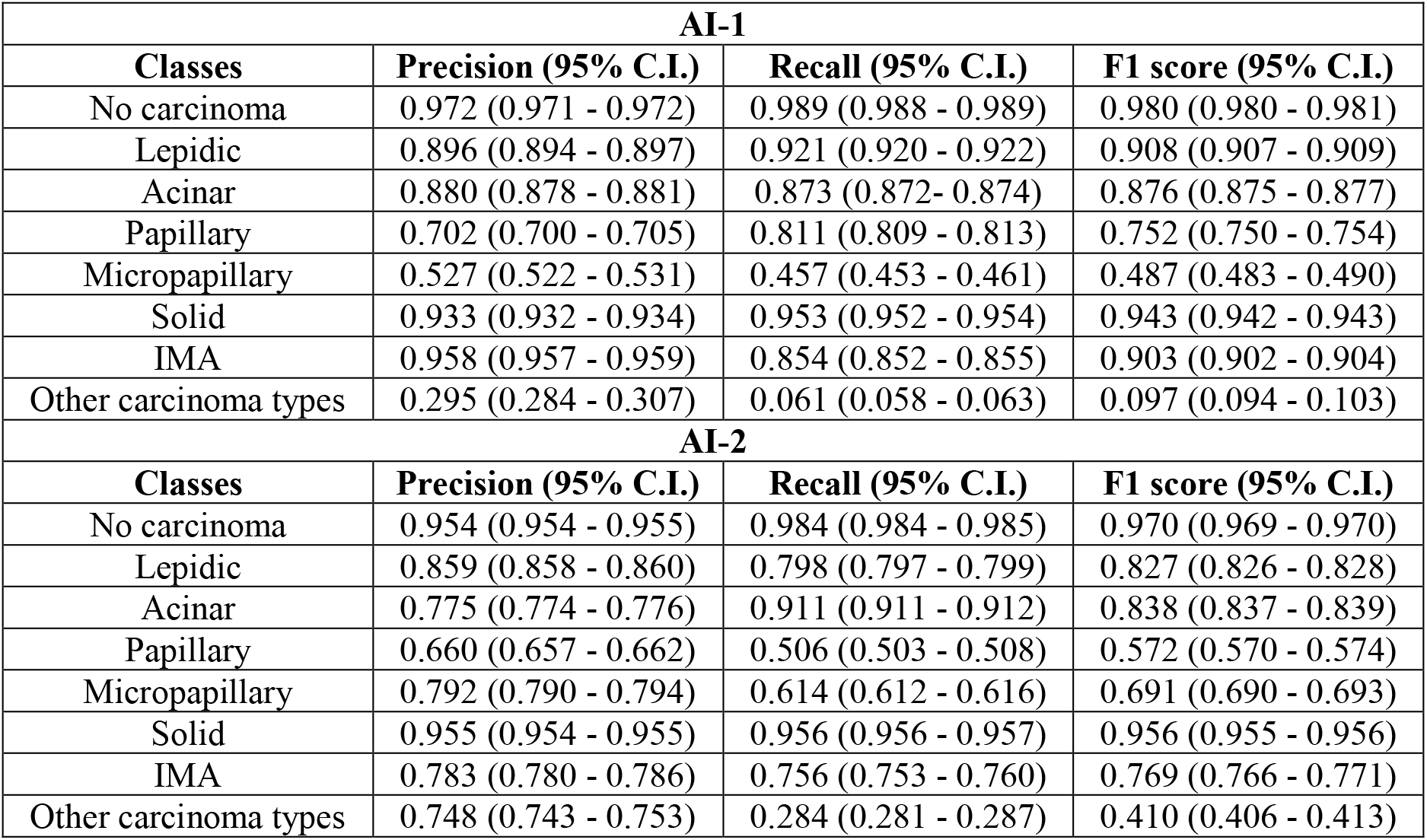
Precision, recall, and F1 score of the trained models per class.

| **AI-1** |  |  |  |
| --- | --- | --- | --- |
| **Classes** | **Precision (95% C.I.)** | **Recall (95% C.I.)** | **F1 score (95% C.I.)** |
| No carcinoma | 0.972 (0.971 - 0.972) | 0.989 (0.988 - 0.989) | 0.980 (0.980 - 0.981) |
| Lepidic | 0.896 (0.894 - 0.897) | 0.921 (0.920 - 0.922) | 0.908 (0.907 - 0.909) |
| Acinar | 0.880 (0.878 - 0.881) | 0.873 (0.872 - 0.874) | 0.876 (0.875 - 0.877) |
| Papillary | 0.702 (0.700 - 0.705) | 0.811 (0.809 - 0.813) | 0.752 (0.750 - 0.754) |
| Micropapillary | 0.527 (0.522 - 0.531) | 0.457 (0.453 - 0.461) | 0.487 (0.483 - 0.490) |
| Solid | 0.933 (0.932 - 0.934) | 0.953 (0.952 - 0.954) | 0.943 (0.942 - 0.943) |
| IMA | 0.958 (0.957 - 0.959) | 0.854 (0.852 - 0.855) | 0.903 (0.902 - 0.904) |
| Other carcinoma types | 0.295 (0.284 - 0.307) | 0.061 (0.058 - 0.063) | 0.097 (0.094 - 0.103) |
| **AI-2** |  |  |  |
| **Classes** | **Precision (95% C.I.)** | **Recall (95% C.I.)** | **F1 score (95% C.I.)** |
| No carcinoma | 0.954 (0.954 - 0.955) | 0.984 (0.984 - 0.985) | 0.970 (0.969 - 0.970) |
| Lepidic | 0.859 (0.858 - 0.860) | 0.798 (0.797 - 0.799) | 0.827 (0.826 - 0.828) |
| Acinar | 0.775 (0.774 - 0.776) | 0.911 (0.911 - 0.912) | 0.838 (0.837 - 0.839) |
| Papillary | 0.660 (0.657 - 0.662) | 0.506 (0.503 - 0.508) | 0.572 (0.570 - 0.574) |
| Micropapillary | 0.792 (0.790 - 0.794) | 0.614 (0.612 - 0.616) | 0.691 (0.690 - 0.693) |
| Solid | 0.955 (0.954 - 0.955) | 0.956 (0.956 - 0.957) | 0.956 (0.955 - 0.956) |
| IMA | 0.783 (0.780 - 0.786) | 0.756 (0.753 - 0.760) | 0.769 (0.766 - 0.771) |
| Other carcinoma types | 0.748 (0.743 - 0.753) | 0.284 (0.281 - 0.287) | 0.410 (0.406 - 0.413) |

Based on the validation results, the ROC curves for both models were plotted, and the corresponding AUC values were calculated (Figures 4C and 4D). For all the classes, the AUC was greater than 0.9, and the highest values were obtained by the “solid” class for both models (0.997 and 0.998 for AI-1 and AI-2, respectively). As the classes of the two trained models had similar AUCs with no statistical differences, we decided to use both models for testing.

### Determination of the lung adenocarcinoma grades of the test set

From the initial 330 WSIs, 239 WSIs representing 142 LADC cases were used to test the two models. Fifteen pulmonary pathologists provided case-level diagnoses of the 142 cases by indicating the percentage of each LADC subtype in a 5% increment. For testing the two models, each WSI of the set was segmented into tiles, and the two models predicted the class of each tile separately. Further, the tiles with attributed labels were reassembled to form the original WSI (Figure 5), and the percentage of each class was calculated for the two models.

**Figure 5.**
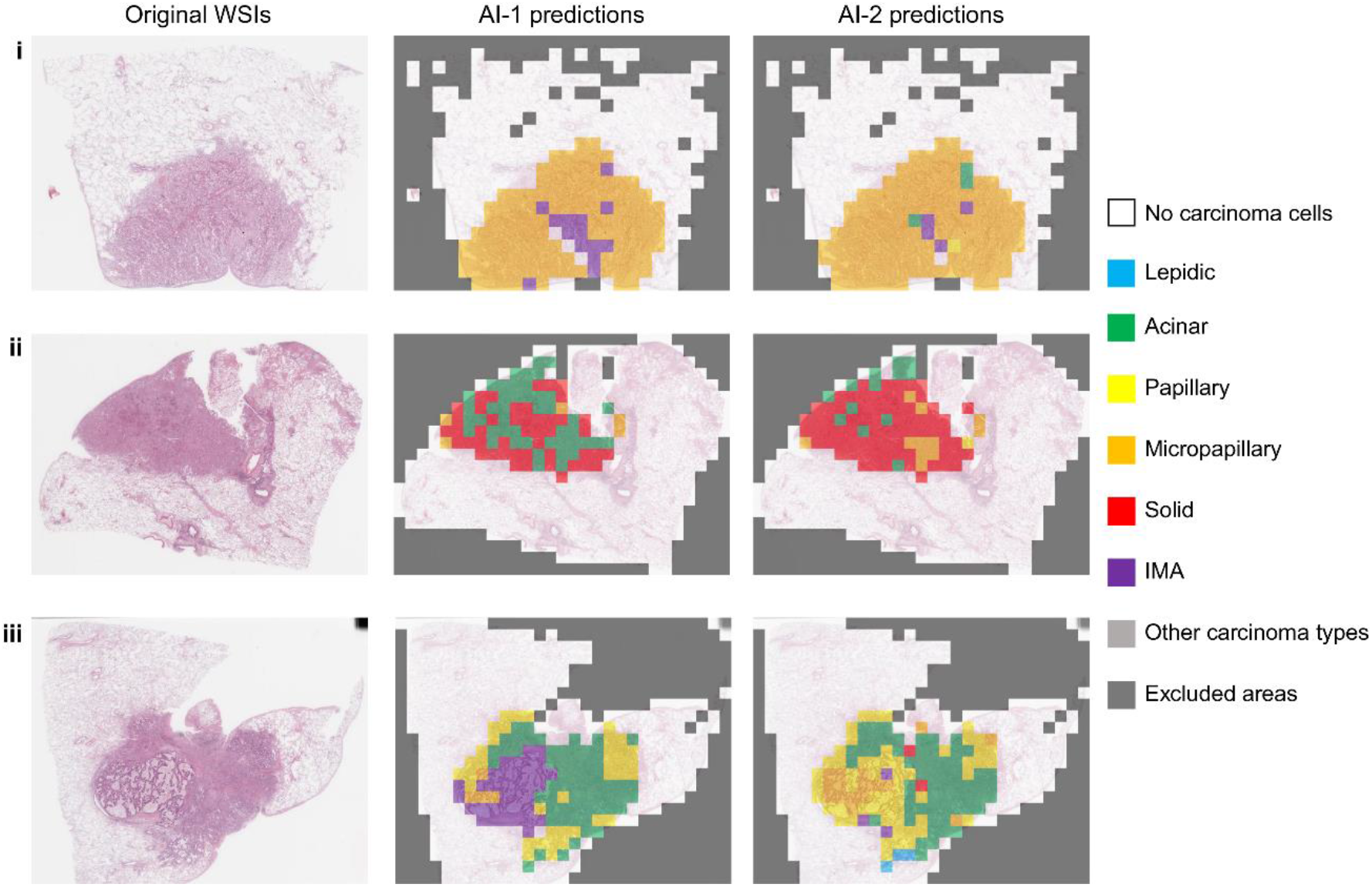
Three example inference maps of the trained models with the predicted lung adenocarcinoma subtypes. Access to the 3 whole slide images is given by the following link: https://bit.1y/3sKXb9M IMA, invasive mucinous adenocarcinoma

The LADC grading [36] was evaluated for each case based on the case-level diagnoses provided by the 15 expert pulmonary pathologists and predicted by the two models (Figure 6 and Supplemental Figure S3A).

**Figure 6.**
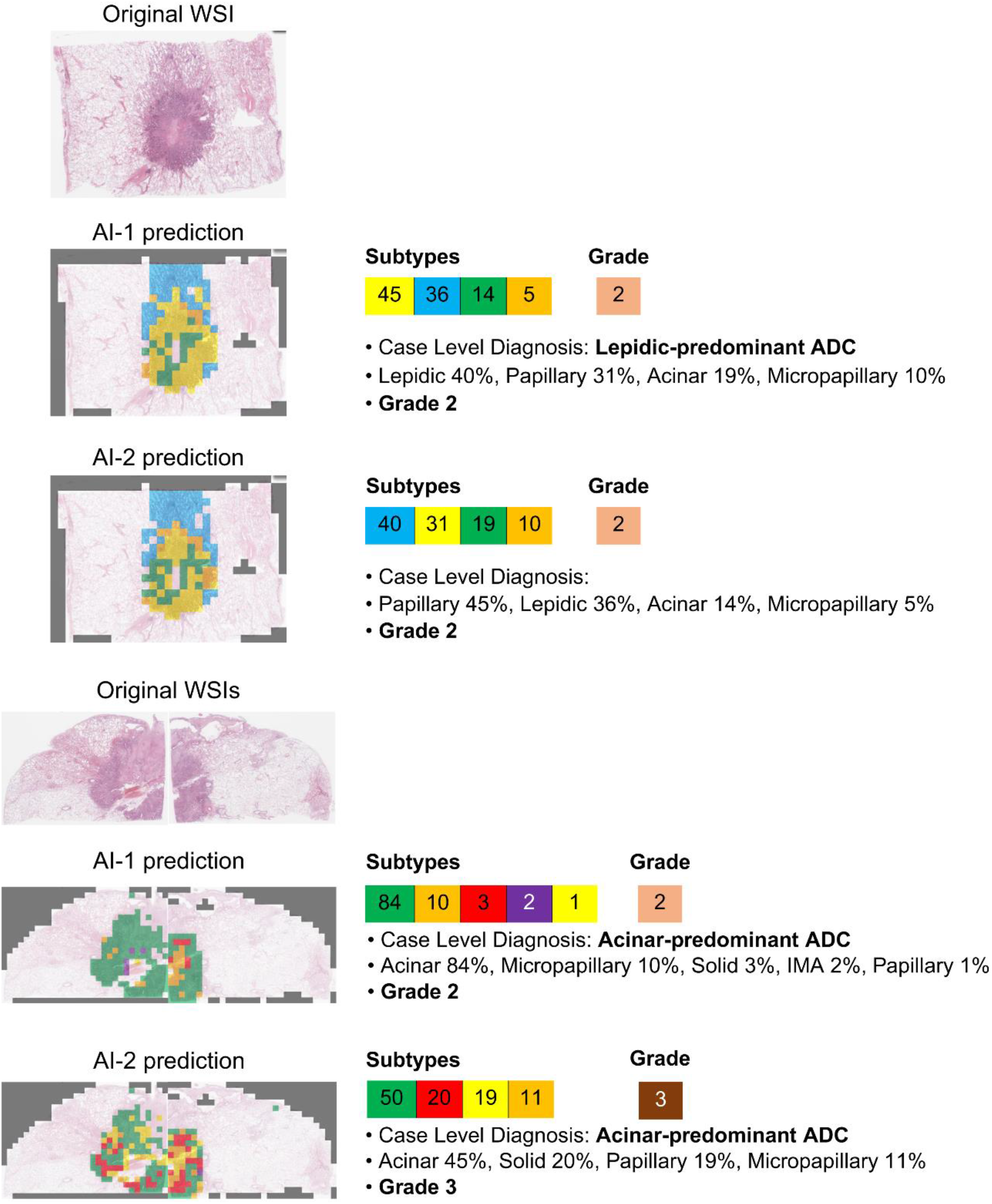
Examples of case-level diagnoses and subsequent IASLC lung adenocarcinoma grades by the two trained models. Access to the whole slide images is given by the following link: https://bit.1y/3sHRFEP ADC, adenocarcinoma, WSI, whole slide image

Excluding cases diagnosed as IMA-predominant by at least one pathologist or one model, the LADC grades were evaluated for a total of 133 cases. The number of grade 1, grade 2, and grade 3 tumours evaluated by each pathologist and model are depicted in supplementary material, Figure S3B. Most of the pathologists and the models assessed the tumours mainly as grade 2, followed by grade 3 and grade 1. The agreement between the pathologist grading evaluation and model grade prediction ranged from 0.314–0.649, which is a slight to moderate agreement. In contrast, the agreement for grading between AI-1 and AI-2 was almost perfect, with a κ value of 0.864 (supplementary material, Figure S3C and Table S1).

### Performance of models on the test set

The LADC grades of the pathologists and deep learning models and the survival data of the 133 patients were used to predict the DSS of the test set. The median follow-up time was 88 weeks for the cohort. Thirty deaths were registered in the cohort, with 130 patients being censored. 102 patients had Stage 0 or IA tumours, 10 patients had stage IB tumours, 14 patients had stage II tumours, and 7 patients had stage III tumours. 68 patients were never smokers. Other patients’ clinicopathological characteristics are provided in our previous study [33].

Clear stratification of the three grades was observed for most pathologists and all the trained models, with a reduction in the DSS as the LADC grades increased (Figure 7). AI-2 (*p*=0.0003) had better separation of the grades than AI-1 (*p*=0.0017). Overall, the two model curves had better statistical significance than most of the pulmonary pathologist curves. AI-2 outperformed every pathologist, except for one (P8) whose grade-driven DSS prediction had the least p-value (*p*=0.0001).

**Figure 7.**
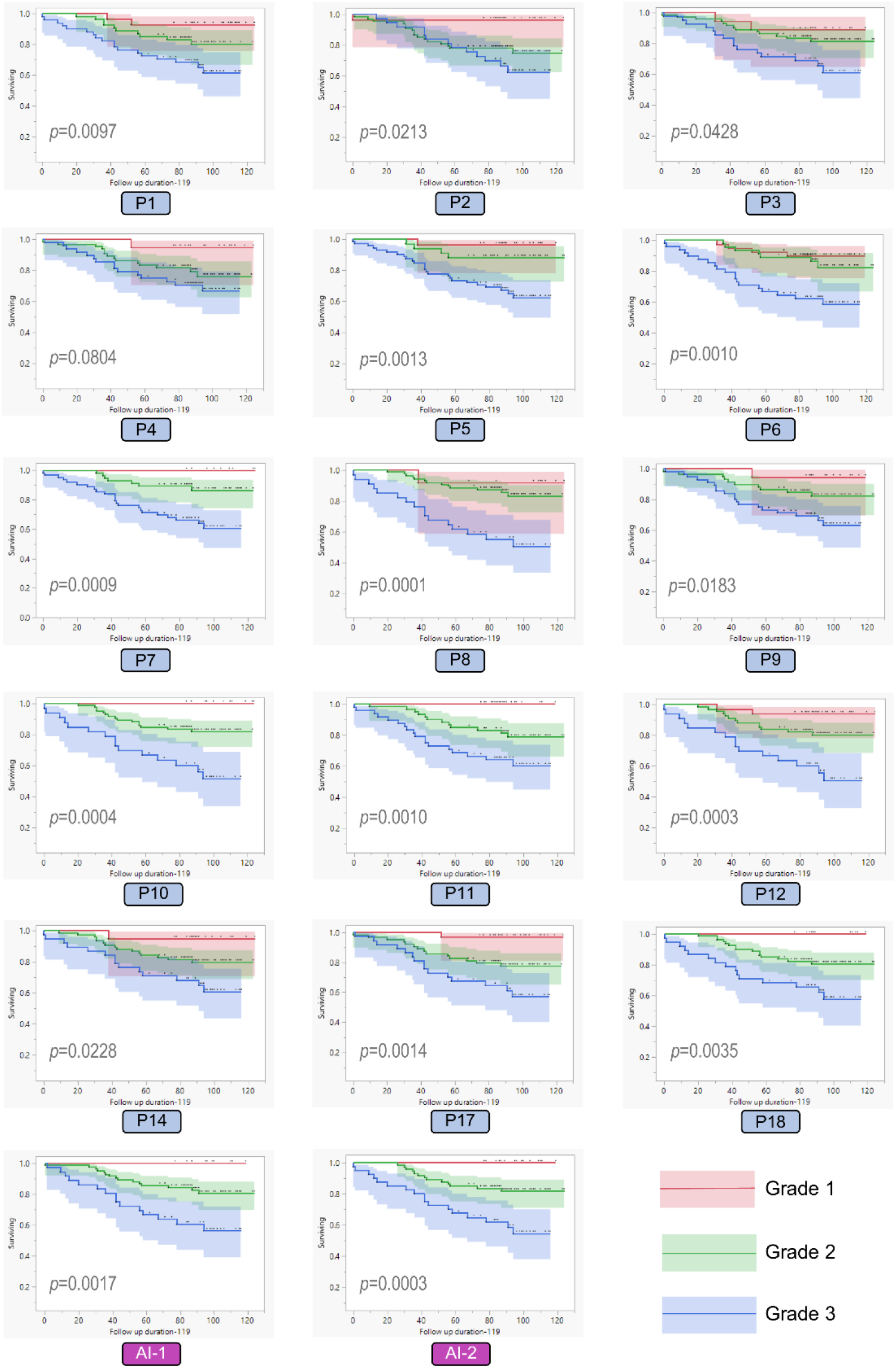
Predicted Kaplan-Meier’s curves of the disease-specific survival of the test set for the lung adenocarcinoma grading system by the 15 pathologists and two trained models P, pathologist

The performance of each pathologist and trained model in case segmentation is represented as the hazard ratio of grade 1 versus grade 2, grade 1 versus grade 3, and grade 2 versus grade 3 in Figures 8A, 8B, and 8C, respectively. Bootstrap sampling was performed 200 times on the grades of the 133 cases of the test set for each evaluator (pathologists and trained models). The ranges of the boxes in the figures show the extent of instability of the segmentation performance of each pathologist or trained model, which varies depending on the samples. As a result, among evaluators, it was observed that both AI-1 and AI-2 exhibited relatively high and stable segmentation performance for each pair of predicted grades as indicated by the relatively high-rank and small variation in the hazard ratio (Figures 8A, 8B, and 8C). Similarly, pathologists P11, P7, P18, and P10 had relatively high ranks with minor variation in the hazard ratio, among the pathologists. In contrast, most pathologists with relatively low ranks (P2, P17, P4, P14, P9, P5, and P8) exhibited less performance stability compared to the trained models in the segmentations of grade 1 and 2, and grade 2 and 3 (Figure 8A and 8C). Although P12, P1, P3, and P6 had stable hazard-ratio variation, their segmentation performance was lower than that of AI-1 and AI-2. The results indicate that the performance of the trained models is high and stable in predicting LADC grades that segment the survival rate, similar or superior to those of expert pulmonary pathologists. Furthermore, trained models provide standardisation of the recognition of the LADC subtypes.

**Figure 8.**
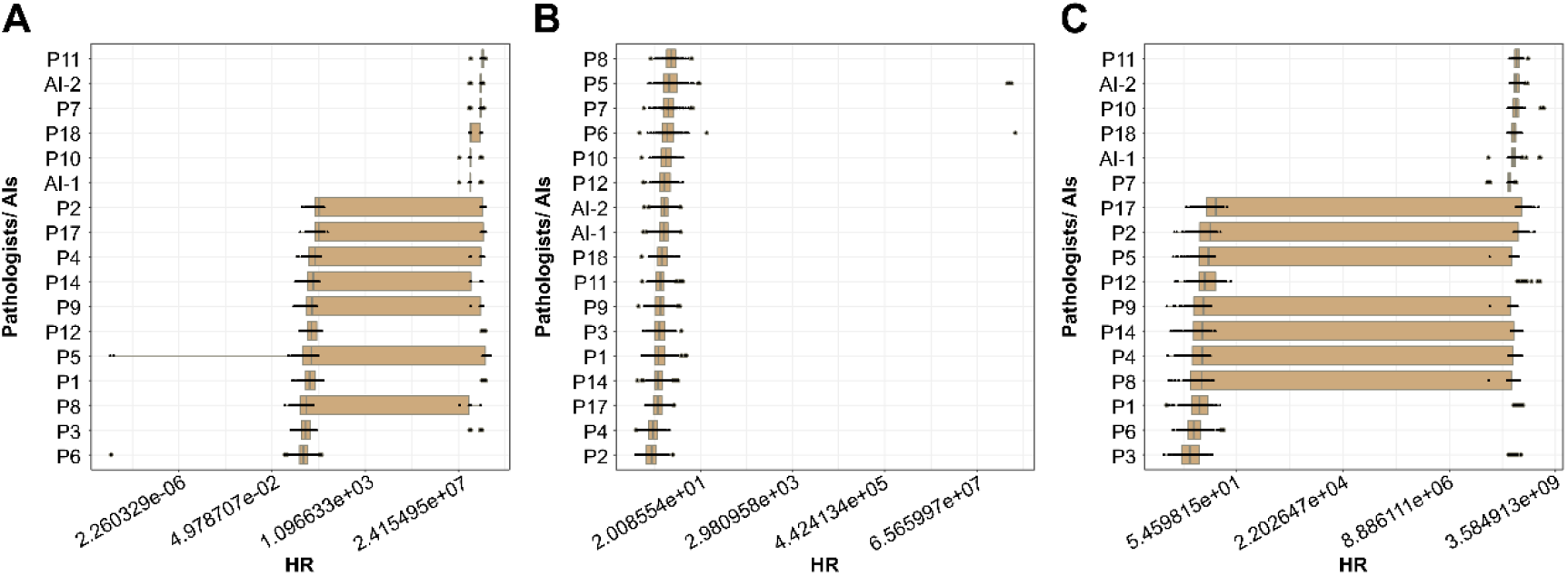
Grading performance of the pathologists and trained models. The box-and-whisker plot shows the 200-times bootstrapped hazard ratio per pathologist and trained model, estimated by the Cox proportional hazard model. The x-axis represents the logarithm of the hazard ratio, and the y-axis represents the pathologists and trained models. The hinges of the box-whisker plot are determined by the 25^th^ percentile and the 75^th^ percentile of the hazard ratios of 200 bootstrap samplings. The median of the 200 hazard ratios is denoted by a grey line in each box. The endpoints of the whiskers show the data points; the minimum and maximum are within a range of 1.5 times the difference between the hinge and median of the hazard ratios of the 200 bootstrap samplings. (A) Hazard ratio between grades 1 and 2. (B) Hazard ratio between grades 1 and 3. (C) Hazard ratio between grades 2 and 3. P, pathologist

## DISCUSSION

This study developed an automated CNN capable of recognising and distinguishing the most common LADC subtypes. Two different sets of consensus LADC images from expert pulmonary pathologists were used to train two models. When tested on an independent cohort of LADC cases, one of the models showed more significant DSS stratification of the three LADC grades than most pathologists and was only slightly outperformed by one of the expert pulmonary pathologists. Moreover, both trained models exhibited low variation in the prediction when evaluating the hazard ratio between different LADC grades. To the best of our knowledge, this is the first study to predict survival based on the LADC grading system using a CNN. The clustering approach used to obtain ground truth images for AI-1 and AI-2 training can also be transposed to other organs and systems with high interobserver variability in the histopathological diagnoses, to develop reliable CNNs. The developed algorithm can assist pathologists in improving the accuracy and in standardising LADC diagnosis. In our previous study [33], we focused on obtaining a reliable set of LADC subtypes ground truth to tackle the interobserver variability. Using a clustering approach, we acquired two different datasets of consensus images from the evaluation of expert pulmonary pathologists. These consensus images were evaluated and showed a reasonable assessment of the prediction of invasive morphology. We, therefore, used these two consensus-image datasets to train two distinct algorithms in this study, AI-1, and AI-2.

The overall precision, recall, and F1-scores of the trained models were high in predicting the LADC subtypes. The highest values were observed for the “no carcinoma”, “lepidic”, “solid”, and “IMA” classes in the two trained models; the highest agreements between pathologists within the same clusters were for the same classes, as shown in our previous study [33]. The results obtained are reinforcing those of published studies focusing on LADC subtyping using deep learning models. Gertych et al. developed CNNs with which the highest F1 scores were obtained for solid tumour growth and non-tumour tissue [34]. On the other hand, the automated classification model developed by Wei et al. reached the best precision, recall, and F1-scores for solid and benign histologic patterns [35]. In contrast, “other carcinoma types” in our study had the least precision, recall, and F1-score among the classes predicted by the two models. This is due to the fewer “other carcinoma types” tiles used for training the two models, as well as the low agreement observed in the separation of this subtype from the other LADC subtypes [33].

Based on the five-fold cross-validation results, the ROC curves were plotted for the two trained models; the AUCs showed the excellent performance of both models in accurately classifying each LADC subtype according to the pathologists’ ground truth. All the classes predicted by the two models had an AUC of over 0.90, which is equivalent to or superior to those reported previously for some of the subtypes. For developing a CNN model capable of predicting the tumour mutation burden status in LADC digital slides, Sadhwani et al. first developed a model for LADC subtype classification, whose AUCs ranged from 0.78–0.98 [40]. Wei et al. in their aforementioned study reported values ranging from 0.97–0.997 [35]. Our models have the advantage of recognising subtypes such as the IMA and “other carcinoma types”, including cribriform architecture or some rare subtypes of LADC.

The pathology committee of the IASLC established a novel grading system for invasive pulmonary adenocarcinoma, which takes into account the predominant histologic pattern and the high-grade pattern with a 20% cut-off for the latter. This grading system better predicts patients’ outcomes compared to systems using mitotic counts or nuclear grade [36]. Several validation studies found that the IASLC grading system had prognostic significance in patients with LADC, with a well-stratified recurrence-free survival or OS [41–44]. Therefore, we used this grading system to evaluate the performance of our trained models by comparing the DSS curves derived by the pathologists and the CNNs. AI-2 had better DSS statistical significance than AI-1, surpassing those of 14 out of 15 expert pulmonary pathologists involved in the study. For the particular cohort used in this study, the result showed that the LADC diagnostic accuracy was improved with one of the trained models. In addition, we performed bootstrap sampling 200 times and evaluated the constancy of the trained models in predicting the LADC grades by estimating the hazard ratio between different grades. AI-1 and AI-2 showed a constant minimal variation in the hazard ratio between different grades, in contrast to some of the pathologists with a wide range of values across the bootstrap sampling data. This finding illustrates the intra-observer variability observed in the recognition of the LADC subtypes, as demonstrated in the reproducibility study by Wright et al. [45]. The trained models not only accurately distinguish the LADC subtypes but also suppress the influence of intra-observer variability through constant and standardisable diagnoses. Additional discussion points about the two models are presented in the supplementary material.

This study is the first to evaluate the performance of a CNN in distinguishing LADC subtypes by predicting the survival of a cohort. We believe that this method demonstrates the effectiveness of the trained models in correctly separating the LADC subtypes in association with their aggressiveness. The training sets of our models included all LADC subtypes, obtained from two sets of consensus tiles deriving from a clustering approach established in our previous study, to overcome influences from interobserver variability [33]. This resulted in a robust dataset for training our algorithms, rendering them capable of distinguishing common tumours such as IMA. However, our study has certain limitations: The usage of single-institutional cases scanned with a single scanner for training the models, and their relatively low number (cases recognised as “other carcinoma types”, in particular). The models were trained with 91 WSIs from which 29,346 tiles were obtained for AI-1 and 30,606 tiles for AI-2. The low-number cases issue was addressed through conventional data augmentation, which in our study resulted in more than 10 million tiles for each model, by applying transforms from the Albumentation library and the mixup principle [38,39]. However, we believe that increasing the training set and including more cases of complex glandular tumour growth or other rare LADC subtypes will refine the prediction accuracy of the model. Furthermore, our models were evaluated on only one cohort. Although the DSS stratification of the LADC grades is satisfactory overall, the model performance needs to be evaluated by including patients from multiple institutions, to generalise the models for widespread usage and increase the LADC diagnostic accuracy. Evaluation of the trained models on multiple cohorts will also reveal differentiation, with one model outperforming the other in the segmentation of LADC grades.

In summary, we developed CNN models for distinguishing the LADC subtypes; the ground truth tiles were obtained from the international panel of expert pulmonary pathologists through a clustering approach. Our models achieved high accuracy and AUC for most of the subtypes, and the model performance was confirmed by the significant stratification of the DSS for the LADC grades, superior to those of most pathologists. We believe that the trained models can improve the LADC diagnostic accuracy and assist pathologists in correctly assessing the patient tumour grades for better follow-up.

## Supporting information

Supplementary discussion points

## ACKNOWLEDGEMENTS

The authors would like to express their gratitude to Kei Tanaka from the department of Pathology, Ai Mori, and Chikako Furukawa from the department of Pathology Informatics, both of Nagasaki University for their assistance in making part of the glass slides.

This paper is based on the results obtained from project JPNP20006 commissioned by the New Energy and Industrial Technology Development Organization (NEDO).

## STATEMENT OF AUTHOR CONTRIBUTIONS

K.L. and J.F. conceived the study. K.L. collected and scanned the glass slides. K.L., K.M., Y.K., T.M., T.T., M.O., and T.N. collected clinical information. K.L., R.A., S.B., L.B., A.C., J.C.E., A.T.F., K.I., Y.K., B.T.L., A.M.M., A.C.R., F.S., M.L.S., K.T., A.M.T., T.T., and J.F. evaluated the tiles and WSIs. K.L., N.O., S.Y., H.S., and J.F. performed model training. N.O., S.Y., W.U., and H.S. validated and tested the trained models. K.L. analysed and interpreted the data. K.L., N.O., S.Y., and S.M. performed statistical analyses. K.L. wrote the manuscript. K.L., A.B., H.S., and J.F. revised the manuscript. J.F. supervised all the data. H.S. and J.F. obtained the grant. All the authors approved the final version of the manuscript.

## LIST OF SUPPLEMENTARY MATERIAL ONLINE

Table S1. Agreements of the IASLC lung adenocarcinoma grading system for the testing set’s cases

Figure S1. Example of tile augmentation using pixel-level and spatial-level transforms and the mixup principle

Figure S2. Example of tiles from the training set with various degrees of agreement. *Complete agreement indicates 10 out of 10 identical labels from pathologists of Cluster-1, and 6 out of 6 identical labels from pathologists of Cluster 2. **Consensus indicates at least 6 out of 10 identical labels from pathologists of Cluster-1, and at least 3 out of 5 identical labels from pathologists of Cluster-2. ***No agreement indicates less than 6 identical labels from pathologists of Cluster-1, and less than 3 identical labels from pathologists of Cluster-2

Figure S3. IASLC lung adenocarcinoma grades of the test set. (A) Pathologist and trained models evaluations of the lung adenocarcinoma grade for each case of the test set. The arrow (Case 68) shows a complete agreement of pathologists for the grade (Grade 3). The arrowhead (Case 79) shows a discordant case where there was no agreement between pathologists (Grade 1, 2, or 3). (B) The number of attributed lung adenocarcinoma grades by 15 pathologists and the two trained models for the test-set cases. (C) Cohen’s kappa scores of the agreement of pathologists’ and trained-model lung adenocarcinoma grades of the test set.

P, pathologist

