## Supplementary tables and figures for "Deep learning for histopathological subtyping and grading of lung adenocarcinoma"

**Supplementary Table S1. Agreements of the IASLC lung adenocarcinoma grading system for the testing set's cases**

(<0 = no agreement; 0–0.20 = slight agreement; 0.21–0.40 = fair agreement; 0.41–0.60 = moderate agreement; 0.61–0.80 = substantial agreement; and 0.81–1.0 = almost perfect agreement). P, Pathologist

| | AI-1 | AI-2 |
|---|---|---|
| P1 | 0.466 | 0.474 |
| P2 | 0.353 | 0.4 |
| P3 | 0.512 | 0.508 |
| P4 | 0.451 | 0.314 |
| P5 | 0.407 | 0.447 |
| P6 | 0.518 | 0.542 |
| P7 | 0.432 | 0.431 |
| P8 | 0.601 | 0.573 |
| P9 | 0.569 | 0.554 |
| P10 | 0.649 | 0.59 |
| P11 | 0.523 | 0.522 |
| P12 | 0.513 | 0.529 |
| P14 | 0.588 | 0.634 |
| P17 | 0.484 | 0.449 |
| P18 | 0.477 | 0.479 |
| AI-1 | | 0.864 |

Figure S1

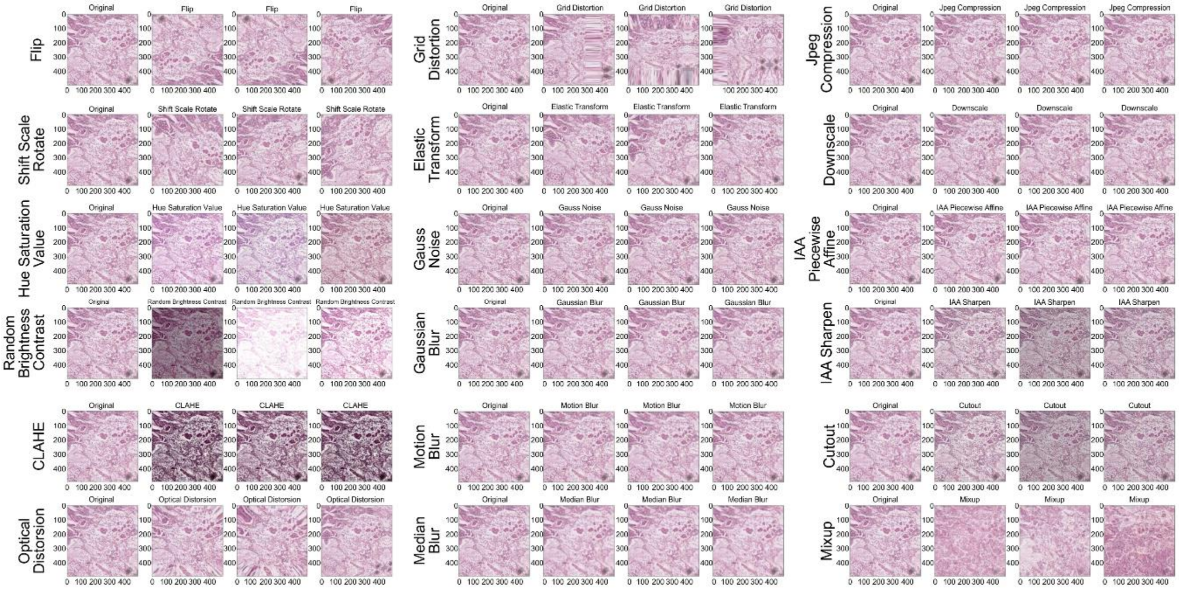


Example of tile augmentation using pixel-level and spatial-level transforms and the mixup principle.

Figure S2

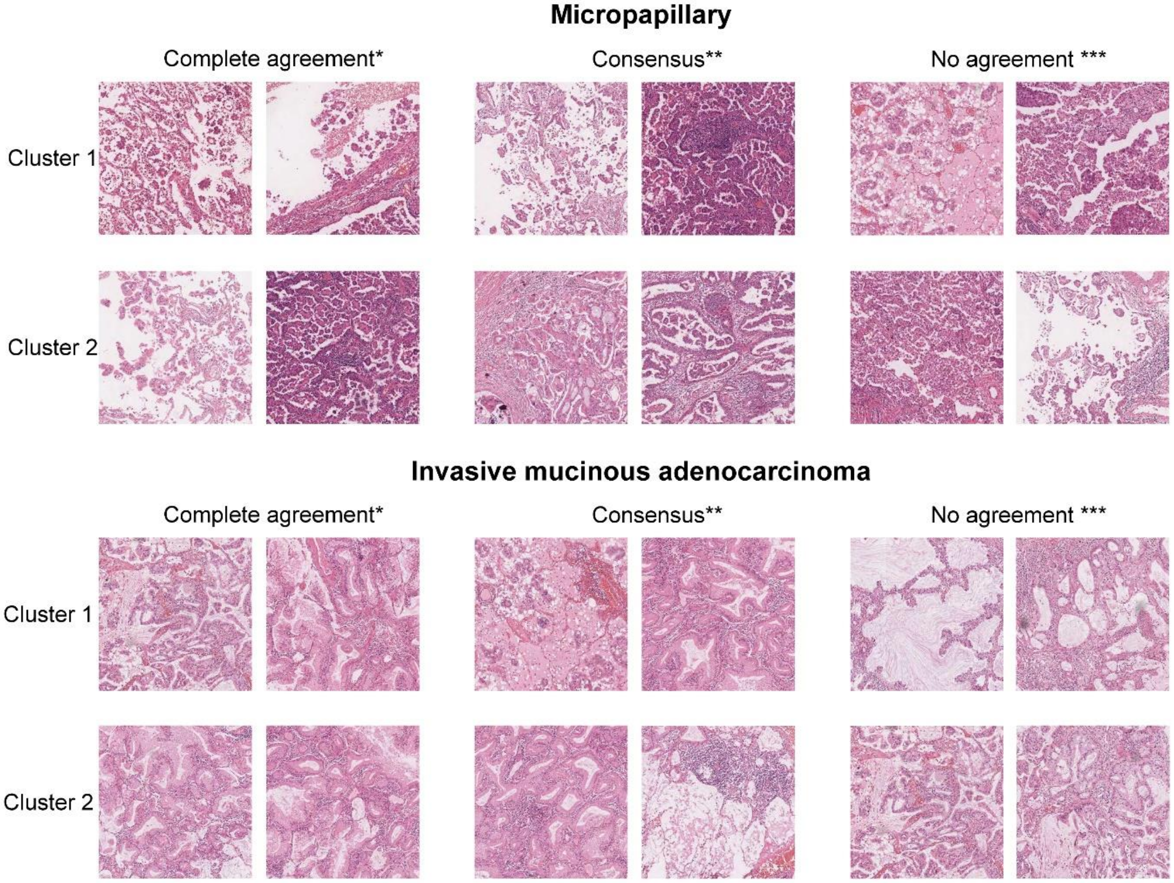


Example of tiles from the training set with various degrees of agreement. *Complete agreement indicates 10 out of 10 identical labels from pathologists of Cluster-1, and 6 out of 6 identical labels from pathologists of Cluster 2. **Consensus indicates at least 6 out of 10 identical labels from pathologists of Cluster-1, and at least 3 out of 5 identical labels from pathologists of Cluster-2. ***No agreement indicates less than 6 identical labels from pathologists of Cluster-1, and less than 3 identical labels from pathologists of Cluster-2

Figure S3

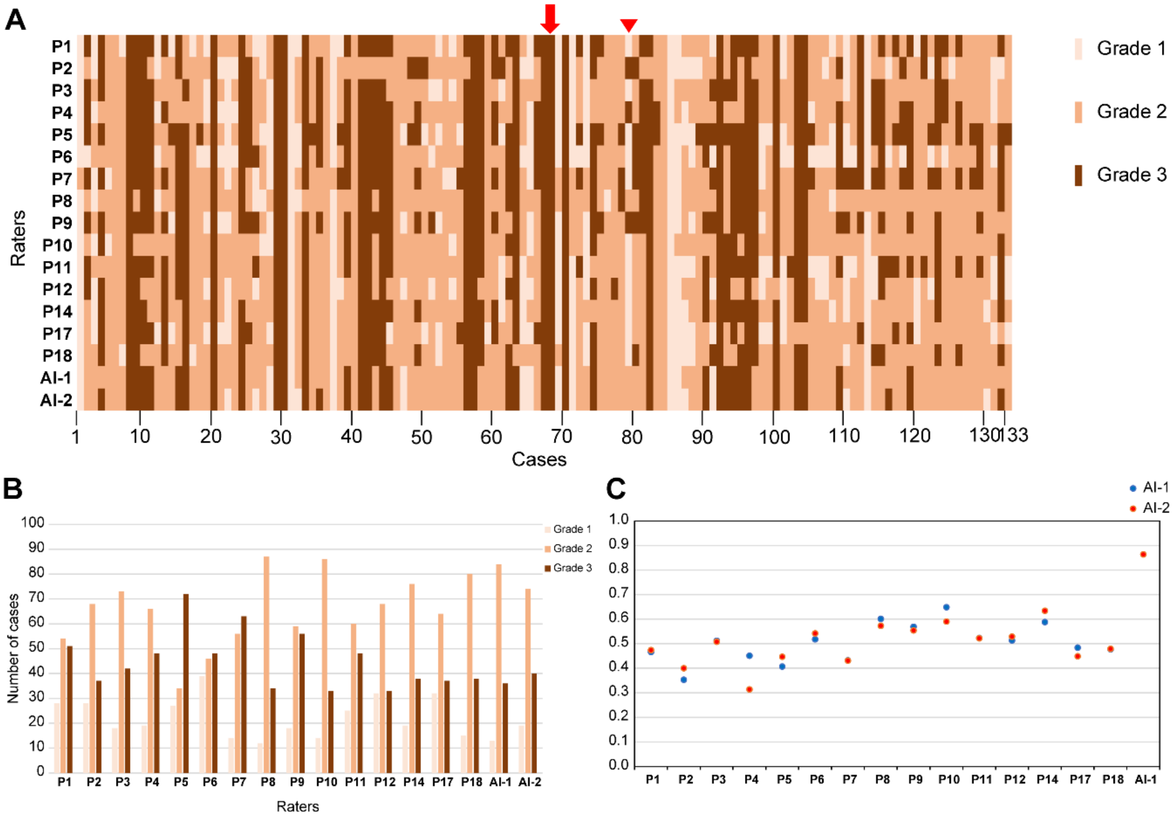


Figure S3. IASLC lung adenocarcinoma grades of the test set. (A) Pathologist and trained models evaluations of the lung adenocarcinoma grade for each case of the test set. The arrow (Case 68) shows a complete agreement of pathologists for the grade (Grade 3). The arrowhead (Case 79) shows a discordant case where there was no agreement between pathologists (Grade 1, 2, or 3). (B) The number of attributed lung adenocarcinoma grades by 15 pathologists and the two trained models for the test-set cases. (C) Cohen's kappa scores of the agreement of pathologists´ and trained-model lung adenocarcinoma grades of the test set.

P, pathologist
