## Supplementary discussion points for "Deep learning for histopathological subtyping and grading of lung adenocarcinoma"

Lami K et al.

### Supplementary Discussion

The two models (AI-1 and AI-2) were trained using consensus images retrieved from the two clusters of pathologists described in our previous study [1]. The main difference between the two clusters was the assessment of a certain pattern as invasive mucinous adenocarcinoma by Cluster-1, and as micropapillary or other carcinoma types by Cluster-2. We decided to use consensus images from the two clusters separately to train two distinctive models, to further determine the best convolutional neural network (CNN). After the validation and testing on an independent dataset, results revealed that both models achieved significance and showed minimal variations of hazard ratio in the segmentation of the lung adenocarcinoma (LADC) grades, with no significant difference between the two. The main aim of the study being the development of CNN to help pathologists in subtyping and grading of LADC cases, it would be practical to only have one model to perform this task. In order to pick up only the best model out of AI-1 and AI-2, additional independent cohorts from multiple institutions will be needed. With the increase of the dataset, we are confident that a difference in the performance will begin to be highlighted between the two models, one outperforming the other with higher rank and less variation of the hazard ratio. This will be subject of an upcoming study.

Another reason of having two models instead of one is that at this point, there is no evidence on whether eliminating differences between the two models will result in a better unique model.  The contrasting pattern recognition from the two clusters may show different association to some clinical parameters, such as therapeutic effect to some particular treatment. Since somehow several experts recognize patterns differently as a group, the recognition may be associated with some unique biological behavior and genetic signatures [2–4]. Therefore, the two trained models was kept, rather than selecting one as a better survival predicting model.
